# Multiplexed detection of nuclear immediate early gene expression reveals hippocampal neuronal subpopulations that engage in the acquisition and updating of spatial experience

**DOI:** 10.1101/2025.07.06.663345

**Authors:** Thu-Huong Hoang, Denise Manahan-Vaughan

## Abstract

Acquired spatial representations are not static. Each re-exposure to the spatial environment stimulates retrieval of the stored experience followed by information re-encoding, including updating if the environment has changed. It remains unclear if the same neurons are involved in these three events.

Here, we used a multiplexed fluorescence in situ hybridization(FISH) approach that detected ‘time-locked’ nuclear immediate early gene(IEG) expression to identify hippocampal neuronal ensembles that were engaged in the acquisition of a spatial representation, as well as its subsequent stabilization and/or updating. Responses were assessed in distal(dCA1) and proximal CA1(pCA1) of male rats. Homer1a was used to detect neuronal recruitment triggered by novel learning of a holeboard environment(HB). cFos and Arc expression was used to detect ensemble stability and/or expansion, or ensemble remodeling, respectively, that was triggered by animal exposure to the now familiar HB that included novel objects(HBO) 25min after the initial HB exposure.

Novel HB exposure resulted in nuclear Homer1a expression in both dCA1 and pCA1. Subsequent HBO triggered significant cFos and Arc expression only in dCA1. IEG co-labelling (Homer1a/cFos, Homer1a/Arc and Homer1a/cFos/Arc) was also only evident in dCA1, reflecting both re-iteration and remodeling of dCA1, but not pCA1 ensembles.

In sum, we show that the contiguous acquisition and updating of spatial representations recruits distinct populations of CA1-neurons reflecting ensemble selection and stabilization, as well as ensemble remodeling. Moreover, whereas dCA1 and pCA1 are involved in the acquisition of the original spatial representation, only dCA1 is engaged in representation updating related to changes in spatial content information.

## 1. Introduction

The acquisition of the initial scaffold of a spatial and/or associative experience can occur in matter of seconds (Piette et al., 2020). However, the acquisition of more detailed spatial representation and/or its updating, is a process that requires time spent in, and movement through, the environment, re-exposure to the same or the updated environment, as well as perception of the saliency and valence of spatial details (Caragea & Manahan-Vaughan, 2021; Griesbauer et al., 2022; Nyberg et al., 2022; Tse et al., 2023). In rats, this process of context- and experience-dependent spatial information acquisition and adaptation is reflected by the stabilization, rate remapping and global remapping of hippocampal place fields (Colgin, 2020) and in the enablement of hippocampal long-term potentiation (LTP) and long-term depression (LTD) by the learning and/or updating of specific components of a spatial representation (Hagena & Manahan-Vaughan, 2024).

Although engram cells have been reported following one-trial aversive learning in rodents (Kupke & Oliveira, 2025), the reactivation of which leads to reinstatement behavioral indicators of memory retrieval (Vasudevan et al., 2024), less is known about the putative role of engram cells in non-aversive forms of spatial learning. Moreover, it has been proposed that rather than have designated neuronal populations that retain specific elements of a spatial representation, the hippocampus utilizes manifold representations of similar (Stacho & Manahan-Vaughan, 2022) or different spatial experiences (Fenton, 2024) that allows the disambiguation, storage and retrieval of context-dependent space.

Subfields of the hippocampus show specialization for the storage, disambiguation and retrieval of different aspects of spatial experience. It has been controversially discussed that different subfields of the cornus ammonis (CA) region of the hippocampus proper, as well as the dentate gyrus, support pattern separation and pattern separation (Kesner, 2013; Quian Quiroga, 2020; Suthana et al., 2021; Yassa & Stark, 2011). However, evidence also exists that the dentate gyrus and proximal CA1 region (pCA1) support pattern separation, whereas the distal CA1 region (dCA1) supports pattern completion (Lee et al., 2020). It has also been proposed that information about non-spatial (e.g. item) identity (‘what’) and allocentric space (‘where’) is delivered to the hippocampus by an ‘offshoot’ of the dorsal and ventral visual streams (de Haan & Cowey, 2011), whereby spatial information is delivered by temporoammonic afferents to pCA1 from the medial entorhinal cortex (EC) and non-spatial information is transferred to dCA1 by lateral EC afferents, resulting corresponding functional compartmentalization of the response of dCA1 and pCA1 to spatial and non-spatial experience (Allison et al., 2023; Vandrey et al., 2021). This interplay was confirmed in studies where nuclear expression of immediate early genes (IEG) was used to study how neurons of dCA1 and pCA1 are engaged in registering novel spatial information: whereas an overt spatial change triggered nuclear IEG expression in both dCA1 and pCA1, the inclusion of novel items into a known spatial environment triggered IEG expression in dCA1(Hoang et al., 2018).

In the present study, our goal was to examine whether differentiated nuclear expression of IEGs resulting from the initial exposure of rats to an overt change in the spatial environment (introduction of a holeboard), followed by insertion of novel physical objects into the holeboard holes, can reveal information encoding and updating dynamics in the dorsal CA1 region of the hippocampus. In particular, we examined the extent to which the same neurons are engaged in information acquisition and updating of a representation of the same spatial environment. Our strategy was based on the latency required for specific IEGs to reach peak nuclear expression after a behavioral learning event (Guzowski et al., 1999; Minatohara et al., 2015). We used fluorescence in situ hybridization (FISH) to detect neuronal Homer1a expression as a biomarker of novel exposure of adult rats to a holeboard. This IEG reaches peak nuclear expression 30-40 min after a specific experience (Bottai et al., 2002; Hoang et al., 2018; Hoang et al., 2021; Hoang & Manahan-Vaughan, 2024; Vazdarjanova et al., 2002). To identify neurons that were engaged in subsequent information encoding and/or updating we used FISH to detect nuclear expression of activity-regulated cytoskeleton-associated protein (Arc) and cFos, both of which show peak nuclear expression 5-6 minutes after a specific experience (Guzowski et al., 1999; Hoang et al., 2018; Saidov et al., 2018; Vazdarjanova et al., 2002). Whereas Homer1a and cFos are transcription factors, Arc is a cytosolic-associated protein (Lyford et al., 1995; Yelhekar et al., 2024). Homer1a plays a role in rendering excitatory synapses amenable for synaptic plasticity (Clifton et al., 2019) and nuclear expression of Homer1a in the hippocampus is increased by induction of long-term potentiation (LTP) and long-term depression (LTD) (Hoang et al., 2021), cellular mechanisms that support spatial learning (Hagena & Manahan-Vaughan, 2024). Associative learning experiences trigger expression of cFos, whereby most studies have examines the role of cFos expressing neuronal ensembles in the acquisition and retrieval of fear memory (Minatohara et al., 2015). A role for Arc in restructuring dendrites and dendritic spines has been described (Chowdhury et al., 2006; Peebles et al., 2010), and several studies suggest that it plays a specific role in the weakening of synapses and/or connectivity within neuronal ensembles (Mikuni et al., 2013; Okuno et al., 2012; Okuno et al., 2018). We thus, used Homer1a as a biomarker of information encoding resulting in the recruitment of neurons into an ensemble, cFos indicated whether these neurons were re-engaged during information updating following object insertion into the holeboard, or whether new neurons were recruited into the ensemble. By contrast, Arc expression was interpreted as indicating the extent to which weaker members of the ensemble were ‘tagged’ for elimination during information updating. Our findings reveal the dynamic nature of ongoing spatial information encoding and indicate that this process is likely to comprise both the strengthening and weakening of hippocampal neuronal networks.

## 2. Methods

### Animals

The study was conducted in accordance with the European Communities Council Directive of September 22nd, 2010. (2010/63/EU) for care of laboratory animals. The experiments were approved in advance by the local state authority (Landesamt für Verbraucherschutz und Ernährung, North Rhein-Westfalia). All efforts were made to minimize the number of animals used for this study, specifically by conducting power calculations to establish the minimal cohort size for meaningful statistical analyses. The animals also served as their own controls. They were housed in sibling groups in a temperature and humidity-controlled vivarium (Scantainer Ventilated Cabinets, Scanbur A/S, Denmark) with a constant 12-h light–dark cycle (lights on from 7 a.m. to 7 p.m.), controlled temperature (22 ± 2 °C) and humidity (55 ± 5%). Food and water were available ad libitum throughout all experiments. In total, 12 male Wistar rats (7–8-week-old) were used for this study. Female rats were not used because adult female rats become stressed by the presence of male rats (DeBold, 1981) and this could alter the outcome of IEG expression and data interpretation. This was also part of our strategy to keep animal numbers to a minimum (reduction of variability of responses) and minimize stressed-related elevations of IEG in the hippocampus (Figure 1A; See, also, comments on FISH strategy, below).

**Figure 1.**
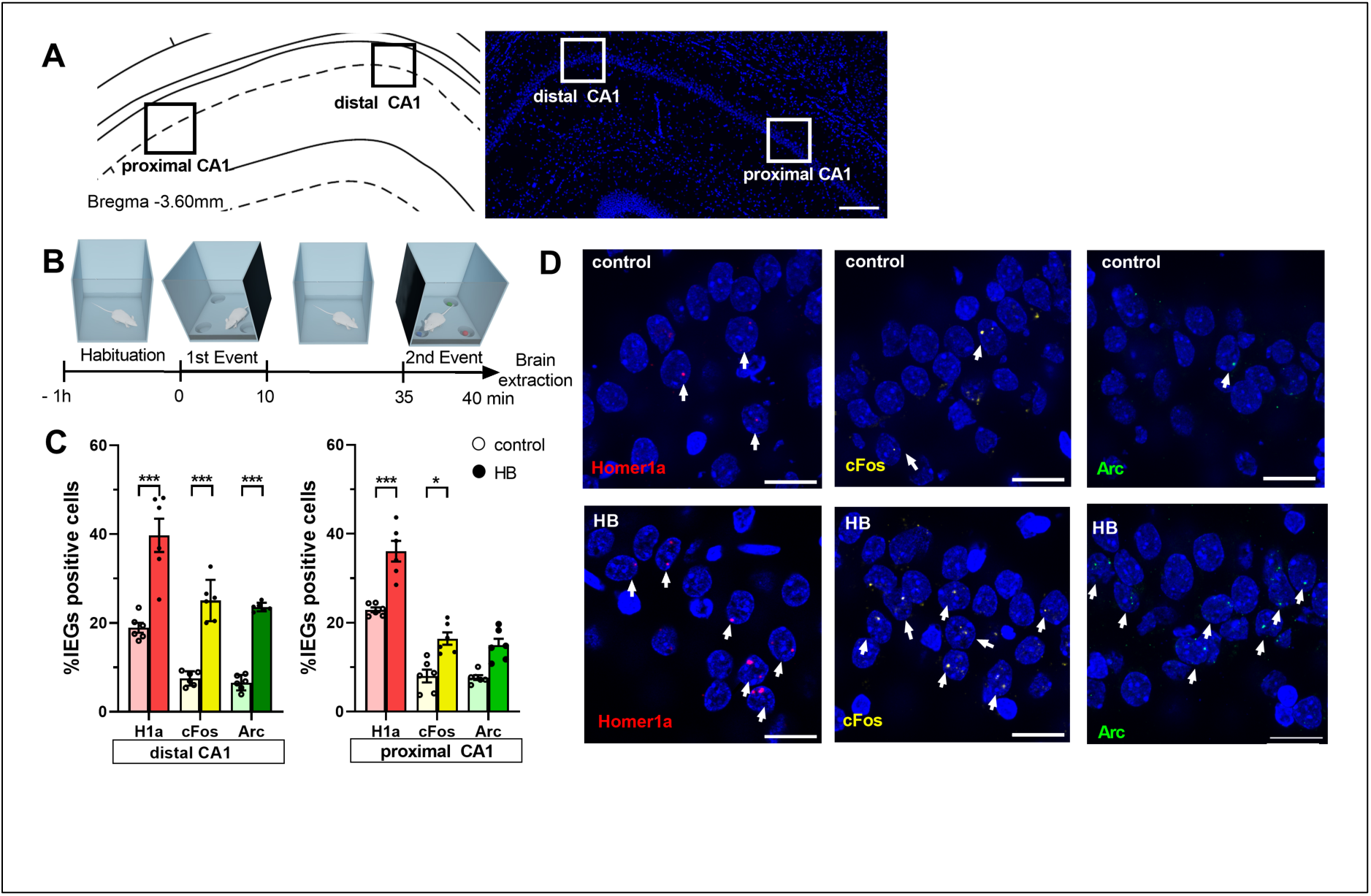
Exposure to novel spatial environment significantly increases nuclear expression of Homer1a, Arc and cFos in the CA1 region. (A) Definition of regions of interest in the CA1 region (left). Image (right) shows a DAPI-stained section at the level of CA1 region of the dorsal hippocampus (3.6 mm posterior from Bregma as determined by a rat brain atlas (Paxinos and Watson, 2014). Z-stacks were obtained in the distal and proximal subcompartments of the CA1 region (black and white squares). Scale bar: 200μm. (B) Experimental design: After having resided in the same experiment room overnight, animals underwent 1h-habituation in the test chamber on the day of the experiment. During the first event, an empty holeboard was introduced to the animal in the chamber. After 10 min exploration time, the holeboard was removed. Animals the rested in the chamber for 25 min. Then, the same holeboard was re-inserted to the recording box, but this time, it contained 3 small objects that were placed inside 3 holes. After 5 min pf exploration, brain extraction occurred. (C) Bar charts show the relative percentage (mean ± SEM) of nuclear Homer1a (red bars), cFos (yellow bars) and Arc (green bars) mRNA expression in neuronal nuclei of the distal (left) and proximal CA1 (right). Novel holeboard (HB) exposure led to a significant increase in Homer1a expression in both distal and proximal CA1 compared to controls (no exploration event). Re-exposure to the holeboard containing novel objects (HBO) significantly triggered cFos expression in the distal and proximal CA1 compared to controls. Significant increases in nuclear Arc expression was only evident in the distal but not the proximal CA1 after HBO (***p<0.001, *p<0.05). (D) Representative images of nuclear Homer1a, cFos and Arc mRNA expression in the distal CA1 of a rat that underwent exploration tasks (HB, bottom row) and in a control rat (control, no exploration event, upper row). Nuclear Homer1a, cFos and Arc mRNA signals are indicated by red, yellow and green dots, respectively. Nuclei were counterstained with DAPI. White arrows indicate IEG positive nuclei. Images were acquired using a wide-field fluorescence microscope at the final magnification of 63x. Scale bars: 20 μm.

**Figure 2.**
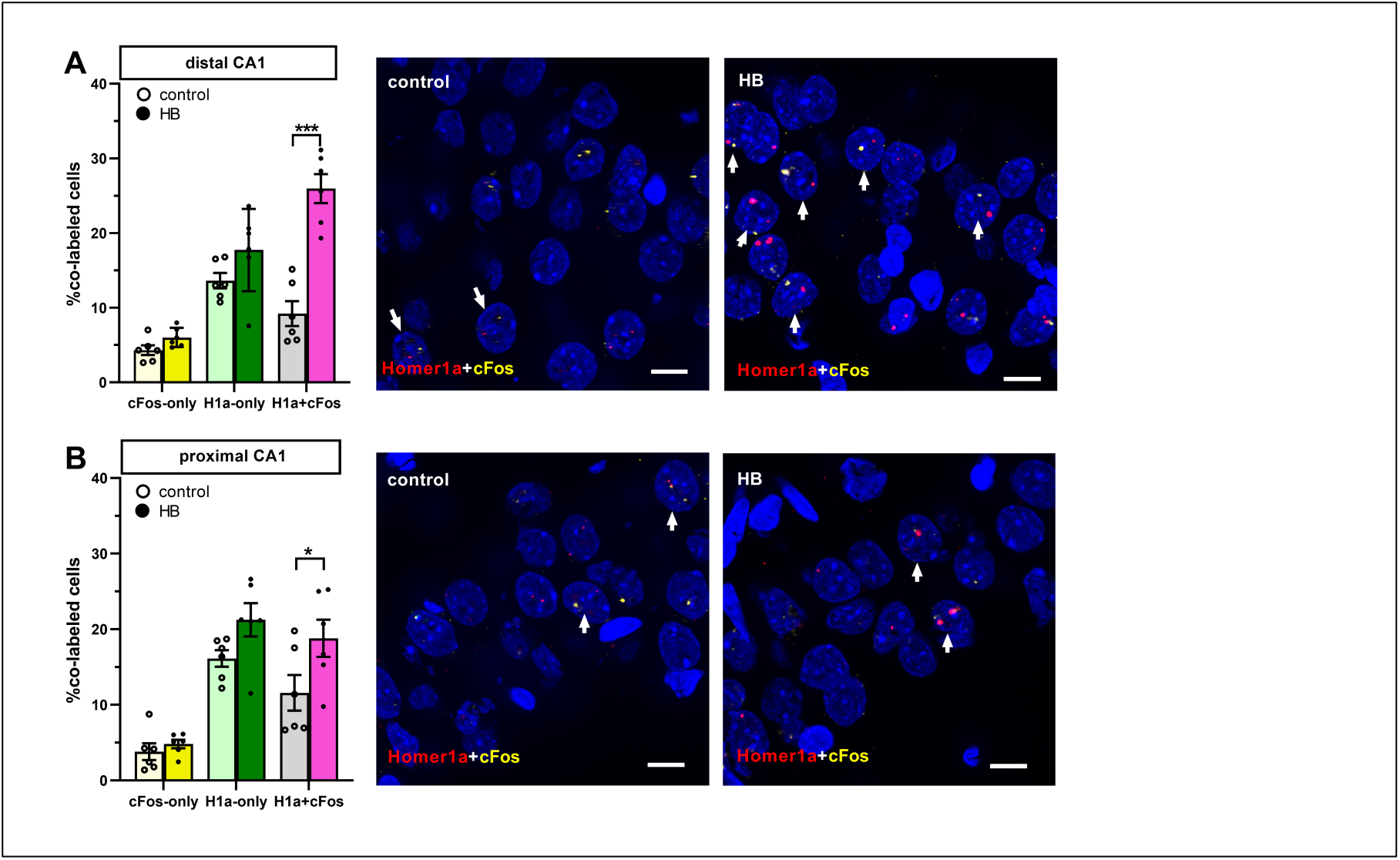
Comparison of neurons co-labeled with Homer1a and cFos reveals ensemble stabilization. **A.** Bar chart (left) shows the relative percentage of neurons in distal CA1 (mean ± SEM) that expressed only Homer1a (dark green, compared to controls light green) following novel HB exposure, or only cFos following HBO (dark yellow, compared to controls, pale yellow). Significant effects were detected in distal CA1when neurons that co-expressed Homer1a and cFos (pink) were compared to controls (gray). (***p<0.001). **B.** A similar comparison of neurons within the proximal CA1 revealed no significant change in Homer1a expression after HB (dark green, compared to controls, light green) or cFos expression after HBO (dark yellow, compared to controls, pale yellow), when neurons that expressed only one of the IEGs were assessed. Significant effects were detected in proximal CA1when neurons that co-expressed Homer1a and cFos (pink) were compared to controls (gray). (*p<0.05). Individual data points are shown as open circles for controls and filled circles for test animals (indicated as HB in chart legend). Right: Representative images of cFos and Homer1a co-labeled neurons in the distal CA1 (upper row) and the proximal CA1 (bottom row) of a rat that underwent exploration events (HB) and in a control rat (no exploration events). Nuclear Homer1a, cFos signals are indicated by red, yellow dots, respectively. Nuclei were counterstained with DAPI. White arrows indicate nuclei that co-expressed Homer1a and cFos. Images were obtained using a wide-field fluorescence microscope at the final magnification 63x. Scale bars: 10 μm.

### Behavioral Experiments

The behavioral habituation and learning protocols were as described previously (Hoang et al., 2018). Animals were handled by the experimenter for 15 min per day for a minimum of 5 days. Then they were habituated to the experiment room and the test chambers for 1h per day, for 3 consecutive days prior to the commencement of the experiments. The test chambers were 40×40× 40 cm in size, open at the top, and were made of gray washable acrylic panels (Perspex®, polymethylmethacrylate, PMMA). The chamber interior was accessible via a translucent PMMA front wall that was held in place by means of tracks installed at the ends of the abutting chamber walls and could be moved upwards by sliding the front wall along these tracks. After the final habituation day, animals were left in the same experiment room overnight. On the day of the first experiment, the animals were placed in the chamber for 1 h before the acquisition phase commenced. The front wall of the chamber was moved upwards so that a holeboard (HB) could be inserted gently into the chamber. The HB was 39.8 x 39.8 x 39.5 in proportions (gray PMMA) and included four holes (with a closed base) that were 5.5 cm in diameter and 5cm deep that were placed equidistantly 2 cm from the edges of HB (Figure 1B).

Behavioral learning events comprised two separate events: The first event comprised exposure to the novel empty HB. Here, the animals hopped onto the HB as it was pushed to touch the back wall of the chamber. The HB was left in place for 10 minutes and then removed (Figure 1B). After HB removal, the animals typically moved to a corner of their chamber and closed their eyes or rested with eyes open.

The second event comprised the re-introduction of the HB into the chamber, whereby this time it included three novel objects (4×2×2 cm) placed (one each) into three of the four HB holes (HBO) (Figure 1B). The total exposure time to HBO was 5 minutes. Whereas novel HB exposure facilitates the expression of hippocampal LTP, novel HBO facilitates LTD in the CA1 region (Kemp & Manahan-Vaughan, 2004, 2008; Manahan-Vaughan & Braunewell, 1999). Events were timed so that HB exposure was aligned with the subsequent detection of peak nuclear Homer1a expression (Hoang et al., 2018) and HBO exposure was aligned with peak expression of Arc (Hoang et al., 2018) and cFos (see below). For this HB and HBO exposure were separated by 25 min (i.e. 10 min HB exposure plus 25 minutes pause), meaning the HBO began 35 minutes after commencing HB. In the interval between HB and HBO, animals resided undisturbed in the recording chamber. The objects used for HBO did not extend above the surface of the holeboard. The animals approached the holes, inserted their noses inside the holes to examine the objects and sometimes lifted them out of the holes with both paws. Interestingly, their typical behavior was to return the objects to the same holes if they had lifted them outside the holes. The configuration of the objects was randomly assigned for each rat.

Immediately afterwards conclusion of HBO (40 min after starting HB exposure) brains were rapidly removed in a smooth movement that involved taking the animals out of the chamber and placing them in a guillotine. This movement took no longer than 3 seconds. Brains were immediately shock frozen in iso-pentane (placed in a small metal container and maintained at a temperature of -60°C), surrounded by dry ice, and then stored at -80°C until further processing. An aged-matched control group of male rats (n=6) was included the study. These animals underwent the same handling and habituation procedures as the test animals. On the day of the experiment, they resided undisturbed in the chamber for at least 1 h. No learning events were implemented. Their brains were quickly removed and flash frozen using the same timeline as described above.

HB and HBO exploration was video-monitored, and the experiment was discontinued, or the data discarded, if an animal spent less than 5 min exploring the empty HB, or less than 3 min exploring during HBO (data not shown).

### Multiplexed Fluorescence In Situ Hybridization

Under RNAse free conditions, brains were sectioned using a cryostat (Leica CM 3050S, Leica Biosystem GmbH, Wetzlar, Germany). Coronal slices (20um) containing hippocampus (from ca. 3.0 to 4.0 mm posterior from Bregma) were collected and mounted directly on superfrost plus ® slides (Gerhard Menzel GmbH, Braunschweig, Germany) and stored at -80°C until further processing.

Arc, Homer1a and cFos cDNA plasmids were prepared commercially (Genscript Biotech, Piscataway Township, New Jersey, USA) using transcripts described by others (Lyford et al. 1995, Brakeman et al. 1997 and Saidov et al. 2019). The cRNA probes were prepared using a transcription kit (Invitrogen Ambion Maxiscript Kit, ThermoFischer Scientific Waltham, USA) and a premixed RNA labeling mix containing Digoxigenin-11-UTP (Roche Diagnostics, Basel, Switzerland) or Fluorescein-12-UTP (Roche Diagnostics, Basel, Switzerland) or Biotin-16-UTP (Roche Diagnostics, Basel, Switzerland). For this study, Arc RNA was labeled with Biotin, Homer1a RNA with Fluorescein and cFos RNA with Digoxigenin. Generated RNA probes were purified using an RNA CleanUp Kit (Monarch®RNA Cleanup Kit, New England BioLabs, Ipswich, USA). Yield and integrity were verified using gel electrophoresis and the concentration was measured by using a QuantiFluor®RNA system (Promega, Madison, USA).

From each animal, we chose 3 consecutive dorsal hippocampal brain sections (ca. 3.6 mm posterior to Bregma) and left the slides at room temperature (RT) until they were defrosted. Later, slides were fixed for 10 min in iced-cold 4% paraformaldehyde in fresh filtered phosphate buffered saline, quickly washed in 2-fold concentrate saline-sodium citrate (2xSSC) buffer (RNAse free), incubated in acetic anhydride solution and briefly washed in 2xSSC (RNAse free) at RT. Then, the slides underwent prehybridization process for 15 min in a mixture of 4xSSC and Formamide (1:1) (RNAse free) at 37°C. The labeled RNA probes were diluted with a concentration of 1g/1ul in 1x hybridization buffer (Sigma-Aldrich, St. Louis, USA), treated at 90°C for 5 min and the quickly put on ice. A humid chamber was prepared with. After the prehybridization, the slides were incubated with the hybridization buffer containing RNA probes for the hybridization process in a humid chamber (with 2xSSC/50 deionized Formamide (Sigma-Aldrich, St. Louis, USA) (1:1) soaked filter paper). The hybridization process occurred in in the humid chamber for approximately 17h at 56°C. Additionally, a negative control FISH test was included to verify the specificity of the hybridized signal. For this, no RNA probes were added to this brain slide (data not shown). After the hybridization, slides underwent stringent washing steps: thrice in 2xSSC at 56°C, then in 2xSSC containing RNAse A at 37°C, again in 2xSSC at 37°C, twice in 0.5xSSC at 56°C, 0.5xSSC at RT, twice in 1xSSC at RT and finally thrice in tris-buffered saline (TBS) at RT. Slides were then treated for 15 min with 3% H_2_O_2_ solution at RT in order to block the endogenous peroxidase.

The Arc-Biotin protocol was previously established (Hoang et al., 2018). Arc-Biotin signal detection was conducted using the following steps:

1. 70 min incubation in TBS-Tween (0.05%, Polysorbate 20) containing 1% Bovine Serum Albumin (BSA) and 20% Streptavidin (Vectorlabs, SP2002, Burlingame, USA).
2. 90 min incubation in TBS-Tween (0.05%) containing 1% BSA and 20% Biotin (Vectorlabs, SP2002, Burlingame, USA) and anti-streptavidin-peroxidase (1:2000, Jackson Immuno Research, Scottsdale, Arizona, USA).
3. Washed 4 times in TBS (5 min each).
4. Incubation for 20 min in TBS containing 1% biotinylated tyramine and 0.01% H_2_O_2_.
5. Washed 4 times in TBS (5 min each) and 60 min incubation in TBS-Tween (0.05%) containing streptavidin CF488 (Biotium, Biotrend GmbH, Cologne, Germany).

The protocol to detect the Homer1a-Fluorescein signal was previously established (Hoang et al., 2018) and comprised the following:

1. Incubation for 70 min in TBS-Tween (0.05%, Polysorbate 20) containing 10% n-Goat serum (Histoprime, Biozol, Hamburg, Germany) and 20% Streptavidin (Vectorlabs, SP2002,).
2. 90 min incubation in TBS-Tween (0.05%) containing 1% n-Goat serum (Histoprime, Biozol, Hamburg, Germany) and 20% Biotin (Vectorlabs, SP2002, Burlingame, USA) and anti-fluorescein-peroxidase (1:2000, Jackson Immuno Research, Scottsdale, Arizona, USA).
3. Washed 4 times (5 min each) in TBS.
4. Incubation for 20 min in TBS containing 1% biotinylated tyramine and 0.01% H_2_O_2_.
5. Washed 4 times (5 min each) in TBS and 60 min incubation in TBS-Tween (0.05%) containing 1% n-Goat serum and streptavidin Cy5 (JacksonImmunoResearch, Scottsdale, Arizona, USA).

The cFos-Digoxigenin signal was detected as follows:

1. Incubation for 70 min in TBS-Tween (0.05%, Polysorbate 20) containing 10% n-Goat serum (Histoprime, Biozol, Hamburg, Germany) and 20% Streptavidin (Vectorlabs, SP2002, Burlingame, USA).
2. Incubation for 90 min in TBS-Tween (0.05%) containing 1% n-Goat serum (Histoprime, Biozol, Hamburg, Germany) and 20% Biotin (Vectorlabs, SP2002, Burlingame, USA) and anti-digoxigenin-peroxidase (1:2000, Jackson Immuno Research, Scottsdale, Arizona, USA).
3. Washed 4 times (5 min each) in TBS.
4. Incubation for 20 min in TBS containing 1% biotinylated tyramine and 0.01% H_2_O_2_.
5. Washed 4 times (5 min each) in TBS followed by 60 min incubation in TBS-Tween (0.05%) containing 1% n-Goat serum and streptavidinCy7 (Invitrogen, Thermo Fischer Scientific, Waltham, Massachusetts, USA).

Later, slides were washed 4 times (5 min each) in TBS, rinsed in double-distilled water, quickly dipped in 70% ethanol and finally stained using 1% Sudan Black B (Merck KGaA, Sigma-Aldrich, St. Louis, USA) in 70% ethanol (Oliveira et al., 2010). Finally, slides were rinsed in distilled water, air dried and mounted in antifading mounting medium (immunoSelect®, Dianova, Hamburg, Germany) containing 4’-6-diamidino-2-phenylindole (DAPI).

Peak nuclear Homer 1a expression occurs 30-40 min after a specific induction event (Bottai et al., 2002). Peak nuclear Arc and cFos expression occur 5-6 min and 6-8min after a specific induction event, respectively (Saidov et al., 2018; Vazdarjanova et al., 2002). Afterwards, the IEG diffuses into the cytoplasm (Bottai et al., 2002; Saidov et al., 2018; Vazdarjanova et al., 2002). For this reason, we assume that if we detect *cytoplasmic* expression of a given IEG at the time-point of nuclear detection, this is an indicator that the animals underwent another salient experience in the time before the experiment was started. Given that the animals resided in their homecages (and subsequently in their homecages) before HB/HBO exposure, this salient experience would have to be stress-related. This can be caused by unexpected noises (in the room or corridor outside the room), strangers entering the room, the presence of adult male rodents (for females), or females in oestrus (for males), or stress-related (22 Hz) vocalisations by animals in the proximity of the room (Cullinan et al., 1995; Inagaki & Ushida, 2017; Manahan-Vaughan, 2017, 2018; Schreiber et al., 1991). For this reason, we ensured that the animals were well-habituated to handling by the experimenter, that only male rats were used and that experiments were conducted under quiet and calm conditions.

### Data analysis

We focused our analysis on the CA1 region, given that this subregion of the hippocampus expresses long-term potentiation (LTP) when novel HB exposure is coupled with afferent stimulation, whereas long -term depression (LTD) is facilitated by novel HBO exposure coupled with afferent stimulation (Kemp & Manahan-Vaughan, 2004). Moreover, novel HB and novel HBO exposure results in a distinct pattern of nuclear IEG expression in the distal and proximal parts of the dorsal CA1 region (Hoang et al., 2018). In other words, the CA1 generates distinct functionally relevant ‘encoding’ responses to novel HB and novel HBO exposure.

Z-stacks were obtained in the distal CA1 (dCA1) and proximal CA1 (pCA1) at a 63x magnification using a Zeiss ApoTome (Fig. 1A). Region of interest (ROI) that contained dCA1 and pCA1 were determined based on their anatomical locations (Naber et al., 2001) and comprised a standardized area of 150μm x 200μm (Fig. 1A). Three consecutive slices of each animal were used for the analysis, whereby we analyzed both hemispheres of each slice and calculated the mean of these three slices. Complete DAPI stained nuclei that showed no evidence of cutting on the edges (of the slides) either in the x, y or z planes, were marked using Fiji software (Schindelin et al., 2012). Neurons were distinguished from glial cells and endothelial cells on the basis of cell morphology and size (Félétou, 2011).

Neurons were checked for nuclear mRNA expression of Homer1a, Arc and cFos that peaked in the nuclei of the CA1 neurons, in an experimenter-blind manner. Based on the fluorescent label, these were detected as red punctae for Homer1a, green punctae for Arc and yellow punctae for cFos (Fig. 1D). Percentages of IEG-mRNA positive cells were calculated per total counted neurons for each subregion of each rat. The designation “positive nuclei” was given to cells that contained intense intranuclear foci of IEG-mRNA fluorescent signals (Fig. 1D). Nuclei that did not contain any intranuclear foci representing a fluorescent signal of IEG-mRNA were counted as negative. The total number of cells analyzed for each ROI of each slide of each animal was on average 80. Then, the relative percentage of single labeled IEG-mRNA positive cells was calculated relative to the total DAPI counts for each ROI. Co-labelling of IEG-mRNA was also assessed. Here, the percentages were calculated by dividing co-labeled counts of IEGs and total IEGs positive counts. Final results are presented as average mean of percentages ± standard error of the mean (SEM) for each group.

### Statistical analysis

Control and test animals comprised n =6 each. All values were verified for normal distribution using the Kolmogorov-Smirnow test with Lilliefors correction. Statistical analysis was performed using Statistica software (version 14.0.1.25, TIBCO Software Inc., Santa Clara, CA, USA). Multifactorial analysis of variance (mANOVA) was performed, followed by a subsequent Tukey HSD post-hoc test for pairwise comparison between factor groups or regions or IEGs. The significance level was set at p<0.05.

## 3. Results

### 3.1. Nuclear Homer1a expression reveals hippocampal neuronal ensembles that encode a novel spatial experience. Responses are distributed accross the distal and proximal CA1

Nuclear Homer1a expression was detected as a biomarker of IEG encoding that was triggered by novel HB exposure. Nuclear Homer1a expression was significantly increased in neurons of both the distal and proximal CA1 region compared to controls (Fig. 1C&D, HB (n = 6) vs. control (n = 6), multifactorial ANOVA for factor “group”: F(1,60) = 221.993, p = 0.0000). A Tukey HSD posthoc test confirmed the significantly greater expression of nuclear Homer1a expression in the CA1 region of the HB group compared to control animals (HB vs. control p = 0.0001 for dCA1, and p = 0.0001 for pCA1). HB-induced nuclear expression of Homer1a was equivalent in dCA1 and pCA1 (Tukey HSD posthoc test for HB, dCA1 vs. pCA1, p = 0.9050). Thus, neurons of the dCA1 and pCA1 respond equally to a novel spatial experience involving a distinct and novel spatial change to the environment.

### 3.2. Nuclear cFos expression reveals that the updating of an established spatial representation requires activation of neuronal ensembles in the distal CA1 region

The IEG cFos is involved in memory acquisition and retention, as shown by many gain-of-function and loss-of-function studies, with responses specifically evident in the CA1 region (Yelhekar et al., 2024). For this reason, we used it as biomarker for ensemble stabilization and/or expansion after novel HBO exposure. Twenty minutes after novel HB exposure, the animals were re-exposed to the HB, but this time novel objects were inserted into three of the four HB holes (HBO). This event was designed to prompt updating of the spatial representation, specifically to include new content information without appreciably changing the spatial environment. The HBO was inserted into the animals’ chamber 25 min after concluding HB exposure and 5 min before brain removal thereby allowing us to use nuclear cFos expression as a biomarker of neuronal activation (Saidov et al., 2018).

Here, nuclear c-Fos expression was elevated in both dCA1 and pCA1 in HBO animals compared to controls (Figure 1C&D, Tukey HSD posthoc test, control vs. HB: p = 0.0001 for dCA1 and p = 0.0256 for pCA1, n = 6 each). This suggests that de novo information encoding occurred as a result of HBO exposure.

To assess whether the same or different neurons responded to HBO compared to HB, we examined co-labeling of Homer1a and cFos (Fig 3). Overall, a higher number of co-labeled positive neurons were evident in the CA1 region compared to controls. (multifactorial ANOVA (F(1,80) = 184.3950, p = 0.0000; control vs. HB: Tukey HSD posthoc test for H1a+c-Fos, p = 0.0001 for dCA1, p = 0.0468 for pCA1, n = 6 each). Interestingly, co-labeling effect was significantly higher in the distal CA1 compared to the proximal CA1 (HB, Tukey HSD posthoc test for dCA1 vs. pCA1: p= 0.0495).

**Figure 3.**
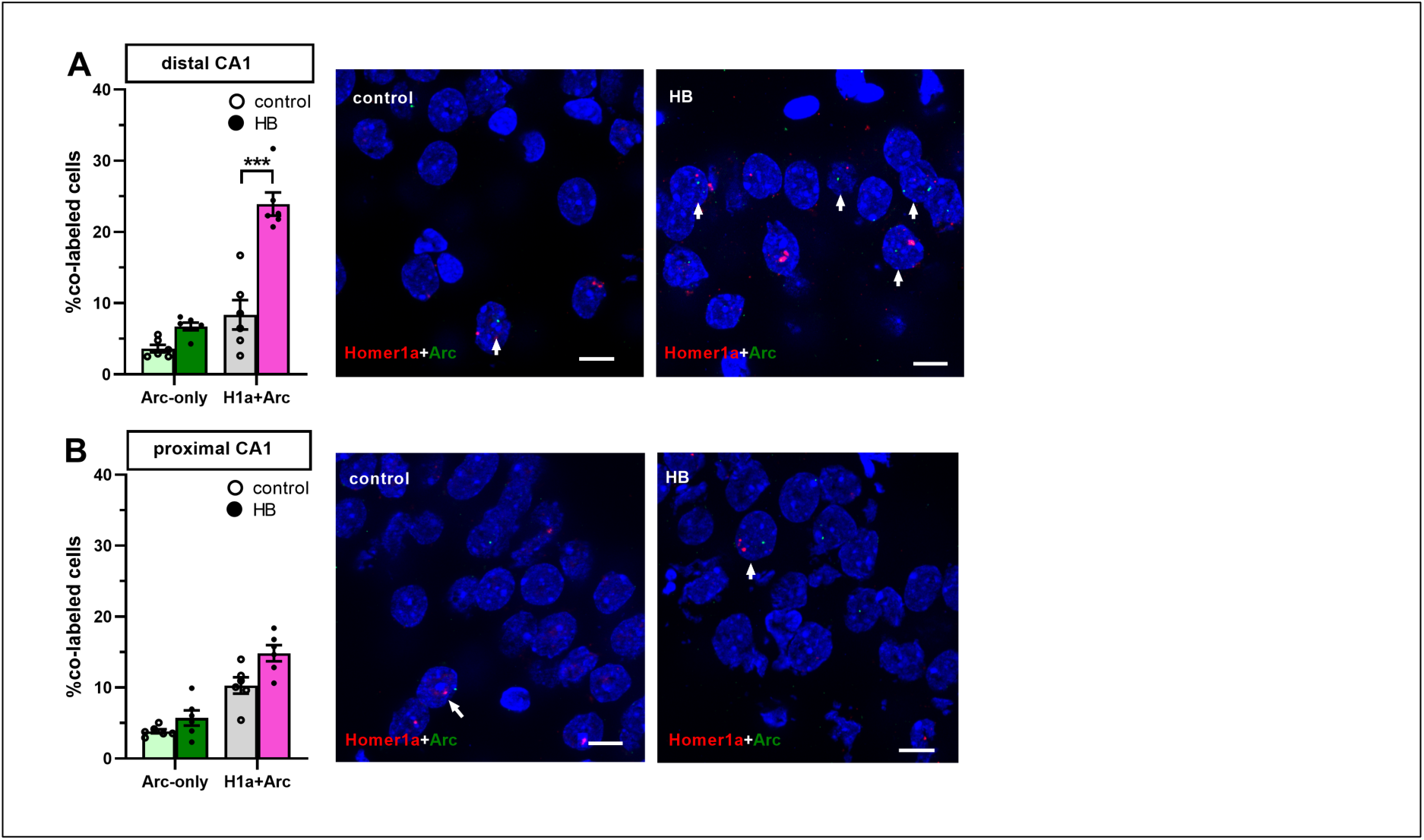
Comparison of neurons co-labeled with Homer1a and Arc reveals ensemble updating. **A, B**. Bar charts show the relative percentage of neurons (mean ± SEM) that expressed only Arc (dark green, compared to controls, light green) following HBO and neurons that co-expressed Homer1a with Arc (pink, compared to controls, gray) in the distal CA1 (**A**) and in the proximal CA1 (**B**) compared to controls (no exploration event). No significant differences were observed in the percentage of neurons of the distal (A) and proximal CA1 (B) that only expressed Arc following HBO, compared to controls. Number of co-labeled neurons was significantly elevated in the distal (A) but not in the proximal CA1 (B) of HB animals compared to control (***p<0.001, *p<0.05). Individual data points are shown as open circles for controls and filled circles for test animals (indicated as HB in chart legend). Right: Representative images of Arc and Homer1a co-labeled neurons in the distal CA1 (upper row) and the proximal CA1 (bottom row) of a rat that underwent novel HB and HBO exploration events (labeled in image as HB) and in a control rat (no exploration events). Nuclear Homer1a, Arc signals are indicated by red and green dots, respectively. Nuclei were counterstained with DAPI. White arrows indicate nuclei that co-expressed Homer1a and Arc. Images were taken using a wide-field fluorescence microscope at the final magnification 63x. Scale bars: 10 μm.

We then examined the extent to which the co-labelled population compared to the populations that expressed Homer1a only, or cFos only (Fig. 3). This was done to clarify whether cFos expression resulting from HBO corresponded to an expansion of the original HB ensemble. No significant difference was detected with regard to cFos-only and Homer1a-only expressing neurons, in both CA1 subcompartments (Tukey HSD posthoc test, control vs. HB: p = 1.0000 for dCA1 and pCA1). However, when we compared this population with the co-labeled neurons, significant effect was observed (ANOVA analysis confirmed that there was a significant difference in these populations (F(1,60) = 39.49, p = 0.0000, n = 6 each).

In dCA1, number of co-labeled neurons was significantly greater than number of neurons that only expressed Homer1a or cFos (HB, Tukey HSD posthoc test: p = 0.0000 for H1a+cFos vs. cFos-only, p = 0.0352 for H1a+cFos vs. H1a-only). In pCA1, neurons that co-labeled Homer1a and cFos were significantly more compared to neurons that only expressed cFos, but not Homer1a (HB, Tukey HSD posthoc test: p = 0.0000 for H1a+cFos vs. cFos-only, p = 0.6777 for H1a+cFos vs. H1a-only). Our results suggest that most of the cells in the dCA1 that were activated by the first event (Homer1a), were reactivated during HBO (cFos). Neurons that were labeled by Homer1a in pCA1 following HB, were not affected by subsequent HBO.

### 3.3. Nuclear Arc expression reveals that the updating of an established spatial representation requires modification of neuronal ensembles in the distal CA1 region

Arc is an IEG that is required for learning and memory (Yelhekar et al., 2024), but in contrast to Homer1a and cFos, it is associated with the elimination of synapses and the remodelling of neuronal ensembles, related to adjustment of synaptic weights (Mikuni et al., 2013; Okuno et al., 2018). We therefore used nuclear Arc-mRNa expression as a biomarker to detect the involvement of CA1 neurons in information updating and ensemble remodelling as a result of HBO exposure following HB exposure.

When we assessed for co-labeling of Homer1a and Arc (Fig. 3), we observed that co-labeling was only significantly present in dCA1 (H1a+Arc, Tukey HSD posthoc test, control vs. HB: p = 0.0001 for dCA1, p = 0.6639 for pCA1). Here, we also compared the co-labeled population with the populations that expressed only Arc and observed a significant effect (HB, Tukey HSD posthoc test, H1a+Arc vs. Arc: p = 0.0000 for dCA1 and p = 0.0012 for pCA1). The significant effect was not observed for the comparison of Arc/Homer1a-labeled neurons, with H1a-only labeled neurons (HB, Tukey HSD posthoc test, H1a+Arc vs. H1a: p = 0.0976 for dCA1 and p = 0.0766 for pCA1, n = 6 each).

Moreover, the number of co-labeled Homer1a and Arc positive neurons was equivalent to the number of co-labeled Homer1a and cFos positive neurons (Tukey HSD posthoc test, H1a+c-Fos vs. H1a+Arc: p = 0.9997 for dCA1, p = 0.8439 for pCA1). This suggests that separate two populations of neurons within dCA1 were engaged in ensemble stabilization (Homer1a/c-Fos) and information updating (Homer1a/ Arc).

### 3.4. Co-labeling of Arc and cFos suggests that active competition between *de novo* encoding and ensemble remodelling contributes to the updating of spatial representations

Although both Arc and c-Fos are required for associative learning, cFos appears to promote the recruitment of neurons into a memory representation (Yelhekar et al., 2024), whereas Arc may be involved in ensemble remodeling and the optimization/updating of established representations (Mikuni et al., 2013; Okuno et al., 2018).

To clarify if HBO resulted in different nuclear expression patterns of these IEGS, we assessed co-labeling in dCA1 and pCA1 after the learning events (Fig. 4). We observed that a significantly greater number of co-labeled Arc and c-Fos positive neurons were detected in dCA1, compared to control animals (Arc+cFos, Tukey HSD posthoc test, control vs. test: p = 0.0001 for dCA1, n = 6 each). By contrast, pCA1 showed no significant differences between two conditions (control vs. test: p = 0.3463). A comparison of the CA1 subcompartments of the animals that participated in the learning events, revealed that co-labeling was significantly greater in dCA1 compared to pCA1 (Tukey HSD posthoc test, p = 0.002).

**Figure 4.**
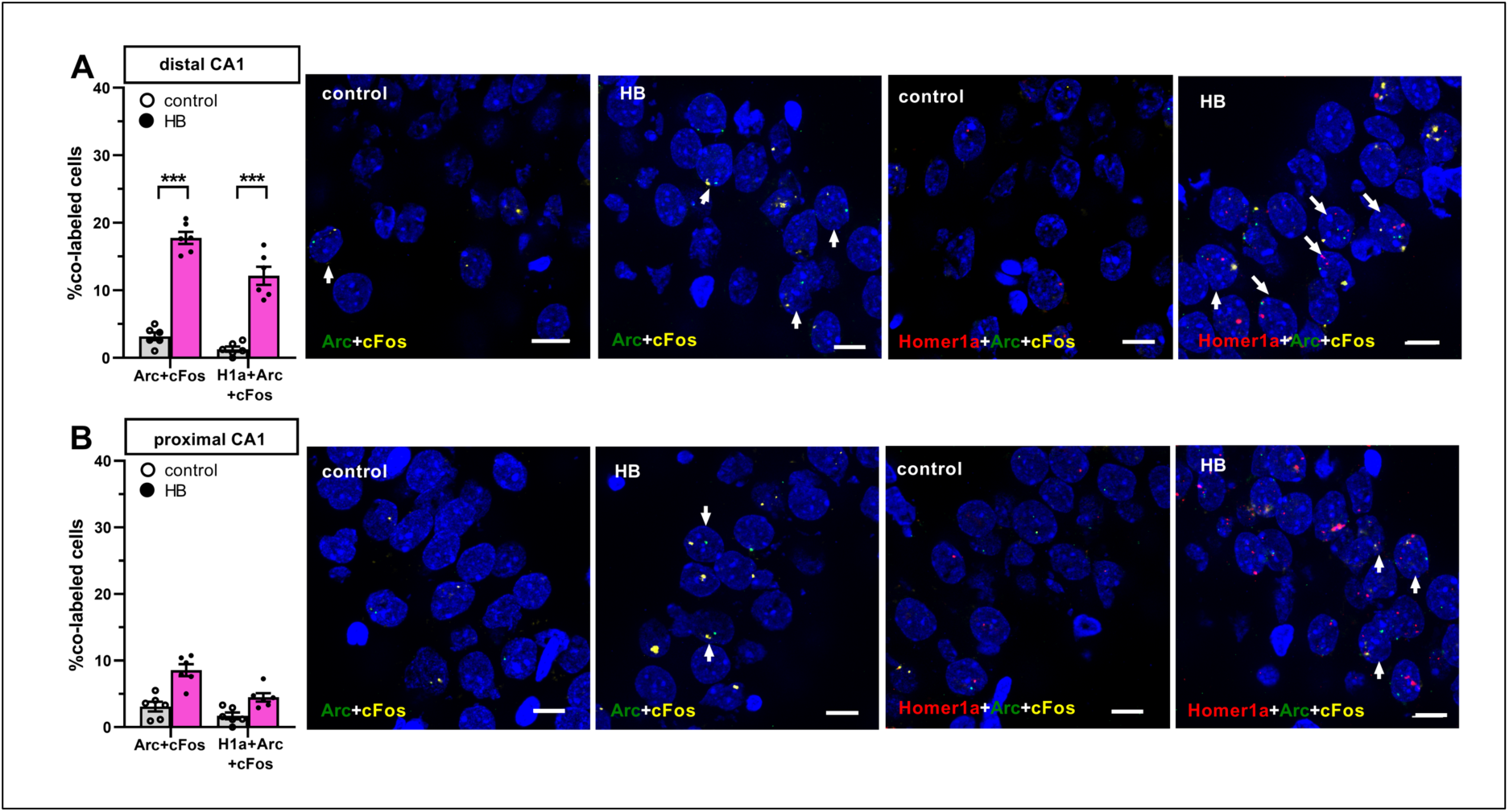
Comparison of neurons co-labeled with cFos and Arc, or cFos, Arc and Homer1a reveals competitive updating of a sparse neuronal population. **A,B**. Bar charts show the relative percentage (mean ± SEM) of neurons that co-expressed Arc and cFos and neurons that co-expressed Homer1a, Arc and cFos in the distal CA1 (A) and in the proximal CA1 (B) compared to controls (no exploration event). The percentage of co-labeled neurons was significantly elevated in the distal (A) but not in the proximal CA1 (B) of animals that engaged in the two learning events compared to controls (***p<0.001). Individual data points are shown as open circles for controls and filled circles for test animals (indicated as HB in chart legend). Right: Representative images of Arc and Homer1a co-labeled neurons in the distal CA1 (upper row) and the proximal CA1 (bottom row) of a rat that underwent exploration events (HB) and in a control rat (no exploration events). Nuclear Homer1a, cFos and Arc signals are indicated by red, yellow and green dots, respectively. Nuclei were counterstained with DAPI. White arrows indicate co-labeled nuclei. Images were obtained at 63x magnification using a wide-field fluorescence microscope. Scale bars: 10 μm.

We then assessed how these populations compared to neurons that expressed cFos alone and Arc alone, as a result of HBO. In dCA1, the proportion of neurons that co-expressed Arc and cFos was significantly greater than the population that expressed Arc alone, or cFos alone (HB, Tukey HSD posthoc test: p = 0.0000 for Arc+cFos vs. Arc-only, p = 0.0000 for Arc+cFos vs. cFos-only). We observed no significant differences in pCA1 when comparing the different populations (HB, Tukey HSD posthoc test: p = 0.3705 for Arc+cFos vs. Arc-only, p = 0.0533 for Arc+cFos vs. cFos-only).

These results indicate that a competition may occur during information updating within a subpopulation of neurons that serves to determine whether the neuron becomes integrated into, or removed from, the updated neuronal ensemble. We hypothesized that those neurons that become successfully integrated into the revised ensemble may co-express all three IEGs.

### 3.5 A sparse population of neurons in the CA1 region exhibit Homer1a, cFos and Arc co-labelling

We detected a small population of neurons in dCA1 that co-expressed Homer1a, Arc and c-Fos signals (Fig 4). Responses were statistically significant compared to controls (Tukey HSD posthoc test H1a+Arc+c-Fos, control vs. test: p = 0.0002 for dCA1, n = 6). No significant differences were observed in pCA1 w (control vs. test: p = 0.9911). Responses were significantly greater in the dCA1 compared to the pCA1 (Tukey HSD posthoc test, p = 0.02). This population of neurons was very sparse compared to other populations that co-expressed only two different IEGs signals (Tukey HSD posthoc test, H1a+Arc+c-Fos vs. H1a+c-Fos: p = 0.0001 for dCA1 and pCA1; H1a+Arc+c-Fos vs. H1a+Arc: p = 0.0001 for dCA1 and p = 0.0003 for pCA1, n = 6 each).

## 4. Discussion

It is a common assumption that neuronal ensembles that are activated by a learning experience, comprise the repository of the acquired memory, although evidence thus far, has only been provided by studies that address fear memory (Frankland et al., 2024; Guan et al., 2016; Tonegawa et al., 2018). More recently however, it was reported that engram cells that are associated with fear memory, are neither static in their expression, nor function (Ko et al., 2025) and instead of reflecting a detailed memory, these cells may correspond to the retention of a gist of the original experience that can be re-utilized in subsequent learning. Little is known about whether engram cells are engaged in benign forms of learning, or indeed about the extent to which neurons, or neuronal ensembles, are re-engaged or updated during associative forms of learning such as benign spatial learning. Here, we report that elements of a neuronal population of the dorsal CA1 region that was recruited by novel holeboard learning, are re-activated during representational updating, but also that members of this ensemble may be eliminated. Concurrently, new neurons are (sparsely) added to the ensemble. Interestingly, neurons of the proximal (pCA1) and distal CA1 (dCA1) regions were engaged in the initial ensemble selection *as well as* ensemble re-activation/stabilization events (as indicated by Homer1a and cFos expression). By contrast increases in nuclear Arc only occurred in dCA1 following exposure to the holeboard with novel objects. This suggests that engram neurons that are putatively activated during *de novo* spatial learning are more stable in pCA1, consistent with its putative targeting by the ‘where’ (dorsal visual) stream, as postulated by others (Amaral & Witter, 1989; Burke et al., 2011; Hoang et al., 2018). The updating of the neuronal ensemble within the dCA1 following item insertion into the holeboard is consistent with the putative role of the dCA1 as recipient of ‘what’ information from the ventral stream (Amaral & Witter, 1989). Moreover, this study indicates that neurons, that are initially recruited during novel spatial learning, form a scaffold that is subsequently modified and updated during integration of further details of this representation.

In this study, we used a multiplexed FISH approach to differentiate neurons of the hippocampal dorsal CA1 region that engaged in nuclear IEG expression as a result of novel holeboard exposure, from those that expressed IEG due to the updating of this environment by the inclusion of objects in three of the four holeboard holes. Previous studies have shown that these two behavioral learning events are tightly associated with the expression of hippocampal synaptic plasticity in the form of persistent LTP and LTD, respectively (Hagena & Manahan-Vaughan, 2024). We focused specifically on nuclear IEG expression in pCA1 and dCA1 because of their abovementioned putative functions in the processing of item-related (what) and location-related (where) elements of spatial information processing. We were particularly interested in the stability of neurons that were activated by the first learning experience (novel holeboard exposure), given that little is known as to whether engram cells are recruited as a result of non-fearful experience. We used Homer1a as a biomarker of neuronal activation during *de novo* holeboard learning, and timed brain removal for FISH assessments to align with the peak nuclear expression of Homer1a after initiation of this experience (35-40 min), (Hoang et al., 2018; Hoang et al., 2021). The specific of this timing was validated by the very low level of cytoplasmic Homer1a staining in our preparations (Figures 1, 2). Animals were well-habituated to the experimenter and test chamber and were kept in a quiet environment prior to experiments. An extensive cytoplasmic signal in the brain slices would reflect prior experience-dependent induction of Homer1a expression, that in our experimental environment could only have been induced by stress. Moreover, the very low cytoplasmic signal confirms that peak nuclear expression of Homer1a occurred 35-40 min after novel holeboard exposure.

We observed that neurons of both the pCA1 and dCA1 showed significant increases in nuclear expression of Homer1, as a result of novel holeboard exposure. Previously, we have reported that the facilitation of hippocampal LTP in the CA1 region of freely behaving rats, by novel holeboard exposure, results in increased nuclear Homer1a expression in the dCA1 and pCA1 (Hoang et al., 2021), as does exposure to a novel holeboard (HB) in the absence of electrophysiological stimulation (Hoang et al., 2018). Effects are consistent with the encoding of both ‘what’ and ‘where’ elements of this spatial experience. We used nuclear Arc and cFos expression as biomarkers of neuronal activation that occurred when items were inserted in the holeboard holes (HBO), this event occurred 20 minutes after novel holeboard exposure and 5 minutes prior to brain removal, thereby allowing us to compare IEG responses with Homer1a expression. Peak nuclear expression of Homer1a occurs ca. 40 min after a specific experience, whereas nuclear Arc and cFos expression occurs 5-6 min and 6-8 min, respectively (Hoang et al., 2018; Saidov et al., 2018; Vazdarjanova et al., 2002). The timing of the experiment (5 minutes of HBO prior to brain removal) plus the time needed for brain removal, will have meant that we were in the range of peak nuclear expression of cFos, as well as Arc. Interestingly, as was the case for Homer1a, nuclear cFos expression was increased in both dCA1 and pCA1 following HB, compared to controls.

This raised the question as to whether this reflected the recruitment of additional neurons into the ensemble, or if the increased cFos expression corresponded to the activation of same neurons that were activated by the *de novo* holeboard exposure. For this we looked for double-labeling of nuclei with cFos and Homer1a. Strikingly, the number of neurons that expressed *only* cFos, or *only* Homer1a was not increased compared to controls. By contrast, the population that co-expressed cFos and Homer1a was significantly greater in both distal and proximal CA1. However, significantly more neurons co-expressed Homer1a and cFos in dCA1 compared to pCA1. This suggests that when animals were re-exposed to the now familiar holeboard that contained novel objects (HBO), the same neurons that were activated during novel holeboard exposure were once more activated during the updating of the holeboard information, especially with regard to ‘what’ information. This finding contradicts reports by others that neuronal cFos expression that is triggered by learning typically overlaps with other IEG-expressing neuronal ensembles (Vazdarjanova & Guzowski, 2004) and rather suggests that the use of benign spatial learning paradigms may reveal nuances of experience encoding that are not possible when fear learning is implemented.

Another possible interpretation of the double-labeling of nuclei with Homer1a and cFos is that this reflected memory retrieval. This is unlikely, however, for the following reasons: Axonal transport of proteins from the nucleus to the synapse takes between 0.2 to 200 mm/day depending on the structure of the protein (Roy, 2020), meaning the fastest transport will take >1 hour, depending on the length of the axon. Homer1a modulates synaptic function, especially through interactions with metabotropic glutamate receptor-5 (mGlu5) (Ango et al., 2000; Chokshi et al., 2019). Given the above-mentioned rates of axonal transport, it is unlikely however, that 35 min after novel holeboard exposure, new proteins, transcribed as a result of nuclear Homer1a expression, would have reached the synapse. Thus, the cFos signal that we detected in neurons that also expressed Homer1a is unlikely to reflect a memory retrieval response and is much more likely to reflect the consistent activation of the same neuron in the novel holeboard and HBO updating events.

Arc plays a specific role in the retention of long-term memory (Attardo et al., 2018; Plath et al., 2006), as well as persistent forms of hippocampal LTP and LTD (Hoang et al., 2021; Messaoudi et al., 2007; Shepherd & Bear, 2011). It has been ascribed a role in the maintenance of changes of synaptic strength within physiological ranges (synaptic scaling) (Park et al., 2008), the spatial distribution of synaptic weights (Okuno et al., 2018), and in the elimination of synapses (Mikuni et al., 2013; Wilkerson et al., 2018). Given that Arc has been reported to support behavioral tagging (Moncada et al., 2011), its role in synaptic scaling and synapse elimination may occur by a process that involves Arc accumulation at activated synapses followed by activity-dependent Arc mRNA degradation (Farris et al., 2014), its role in AMPA receptor endocytosis (Park et al., 2008; Plath et al., 2006). Based on these findings, we interpreted nuclear Arc signals as indicating that a neuron will be removed from the previously created ensemble. Consistent with this interpretation we did not detect an increase in the percentage of neurons that were labeled with Arc alone, compared to controls. However, we detected a significant population of neurons in dCA1 that co-labelled for Homer1a and Arc. Given that dCA1 is believed to process ‘what’ aspects of information, the co-labeling of neurons with Homer1a and Arc could on the one hand reflect the updating of the neuronal ensemble with novel item information, and on the other hand, indicate that co-labeled neurons will be eliminated from the original dCA1 ensemble. If the latter possibility is valid, we should also see a significant percentage of neurons that co-label with cFos and Arc in dCA1. This was indeed the case. A similar significant outcome was obtained when we assessed the percentage of neurons that co-expressed Homer1a, cFos and Arc. Thus, these neurons may have been subsequently removed from the ensemble, or may have been engaged in a competitive process whereby, their synaptic weights were adjusted in a spine-specific manner.

In summary, we show in this study that a neuronal ensemble is activated in the cornus ammonis by *de novo* spatial learning that is distributed across the dCA1 and pCA1 (as reflected by nuclear Homer1a expression), consistent with the integration of information about ‘what’ and ‘where’ aspects of space, respectively. The subsequent introduction of novel objects into the holeboard holes prompted a re-activation of the previously activated dCA1 ensemble (as reflected by co-labeling of nuclei with Homer1a and cFos), presumably related to the integration of new ‘what’ information in the holeboard representation, and re-exposure to the now familiar holeboard. This process was also accompanied by the expression of Arc in the dCA1 of the previously activated ensemble, consistent with an experience-dependent modification of the neuronal ensemble triggered by updating of the holeboard representation. These findings suggest that experience-dependent nuclear IEG expression is not a homogenous phenomenon and that different neurons within an ensemble may encode different information. Moreover, Homer1a may be a useful biomarker of cells that stably retain encoded information, whereas Arc may serve as a biomarker of experience-dependent changes in ensemble dynamics.

## Data availability statement

The raw data supporting the conclusions of this article will be made available by the authors, upon reasonable request.

## Ethics statement

The animal study was approved by the Landesamt für Verbraucherschutz und Ernährung, North Rhein-Westfalia, Germany. The study was conducted in accordance with the local legislation andinstitutional requirements.

## Author contributions

The study was designed by Denise Manahan-Vaughan. Experiments, design of RNA probes and implantation of behavioral experiments and FISH protocol and analysis were conducted by Thu-Huong Hoang. Denise Manahan-Vaughan and Thu-Huong Hoang interpreted the data and wrote the paper.

## Acknowledgements

This work was supported by a grant from the German Research Foundation (Deutsche Forschungsgemeinschaft, DFG) to Denise Manahan-Vaughan (SFB 1280/A04, project number: 316803389). We gratefully acknowledge Ute Neubacher for her advice on triple labeling FISH protocol and Beate Krenzek for her support with preparation of RNA probes and FISH procedures. We thank Juliane Böge for her assistance in behavior experiment and Nadine Kollosch for animal care.

## Generative AI statement

The authors declare that no Gen AI was used in the creation of this manuscript.

